# GC3 codons enhance protein production in diverse GC- and AT-rich plant species

**DOI:** 10.1101/2025.11.25.690434

**Authors:** Isaiah D. Kaufman, Hsin-Yen Larry Wu, Polly Yingshan Hsu

## Abstract

Engineering translation holds great promise for maximizing protein yields in agriculture and biotechnology, but the diversity of plant genomes hinders predictable engineering. To identify mRNA features that broadly improve translation, we conducted a comparative translatome analysis across model plants. We found that codons with G or C at the third position (GC3) are consistently associated with higher translation efficiency. Experimental results confirmed that elevating GC3 increases both protein output and mRNA abundance, in both GC- and AT-rich species. Comparative analyses across 80 plant species, spanning a wide range of GC3 levels, show that GC3 content is positively correlated with translation efficiency. Additionally, high GC3-codon usage is conserved among endogenous high-abundance proteins, such as Rubisco small subunits and ribosomal proteins. Finally, tRNA availability likely explains why GC3 codons broadly enhance translation. Together, our results provide a simple guideline for codon optimization: increasing GC3 can enhance protein production across diverse plants.

## INTRODUCTION

Among the various layers of gene expression regulation, translation represents a promising target for engineering gene expression due to its direct impact on protein production. Manipulating mRNA features that influence translation has enabled the modulation of protein output in diverse plant species ^1–5^. Gene-specific regulatory elements, such as upstream open reading frames (uORF) in 5′ UTRs and binding sites for specific RNA-binding proteins often in 3′ UTRs, provide transcript-specific translational regulation ^6–9^. In contrast, certain sequence features exert more global effects on translation. For example, the nucleotide context surrounding the start codon, known as the Kozak sequence, plays a critical role in translation initiation across eukaryotes ^10–13^. In GC-rich plants, such as rice and maize, manipulating codon usage, specifically the GC content at the third nucleotide position of the codon (GC3), has high potential for enhancing protein production. In rice, polysome profiling revealed that highly translated mRNAs are associated with higher GC3 levels ^14^. In maize, increasing GC3 content in transgenes elevates both mRNA and protein levels, likely by reducing siRNA-mediated gene silencing ^15^, though its role in translation remains to be investigated.

Codon optimization, which substitutes synonymous codons to match the host species’ codon usage, has long been employed for heterologous protein production ^16,17^. The availability or complete lack of certain tRNAs is a major reason why codon optimization is often necessary, particularly for expression in prokaryotic species ^18–20^. In eukaryotes, tRNAs have diversified, increasing both the total number of tRNA genes and the diversity of anticodons ^21^. Additionally, wobble base-pairing allows certain tRNA anticodons to recognize multiple codons ^19,22,23^. While codon optimization has successfully increased target proteins in many plants ^24–27^, it is not universally effective and, in some cases, has led to reduced protein production ^28–30^. These findings suggest that codon usage is subject to additional regulation that remains to be elucidated across plants.

The diversity of crops and plants overall presents a challenge for our ability to reliably predict and engineer translation. Early studies analyzing the annotated coding sequences have shown that for the Kozak sequence, eudicots preferentially use AAA**<u>AUG</u>**GC, whereas monocots prefer GCC**<u>AUG</u>**GC (**<u>AUG</u>** indicates start codon) ^12,31,32^. For codon usage, eudicots preferentially use A and U in the 3^rd^ position of each codon, while monocots prefer G and C ^16,33–35^. Yet, it remains unclear how these sequence features directly affect translation globally, as translation is a more onerous metric to quantify until recently.

Our ability to quantify the translation efficiency (TE) of individual mRNAs has been revolutionized by ribosome profiling (Ribo-seq). Ribo-seq is a method for deep sequencing of ribosome footprints, and can be used to infer TE when normalized to transcript abundance from RNA-seq ^36^. Ribo-seq has revealed 100-fold differences in TE in yeast, and 4000-fold differences in TE in maize ^36,37^. Using hundreds of Ribo-seq datasets in yeast, mammals, and humans, in-depth models have been developed in these species to predict TE of a given mRNA based on its sequence features ^38–42^.

Here we aimed to uncover mRNA features that universally increase translation efficiency and protein production in diverse plant species. We employed both computational and experimental strategies and found that increasing GC3 could be widely applied to enhance protein production in both AT-rich and GC-rich plants. The conserved GC3-favoring tRNA pools help explain why GC3 links to high translation. We propose that increasing GC3 could be a universal codon-optimization guideline for maximizing protein yields in plants and possibly other eukaryotes.

## RESULTS

### Identifying mRNA features associated with high translation efficiency

To identify mRNA features associated with translation efficiency (TE) regulation across plants, we sought to compare mRNA sequences between high-TE genes and low-TE genes from monocots and eudicots. To this end, we analyzed Ribo-seq and RNA-seq data from two monocots, maize (*Zea mays* B73) ^37^ and rice (*Oryza sativa* Nipponbare) ^43^, and two eudicots, Arabidopsis (*Arabidopsis thaliana* Col-0) ^9^ and tomato (*Solanum lycopersicum* Heinz 1706) ^44^. Using the Ribo-seq and RNA-seq ratio, we calculated TE for >17,000 genes per species. Subsequently, we defined genes with the highest and lowest 10% TE as high-TE genes and low-TE genes in each species (**Fig. 1A**). Consistently, protein abundance measured by quantitative proteomics shows that high-TE genes are associated with higher protein levels compared to low-TE genes (**Fig. 1B**).

**Figure 1:**
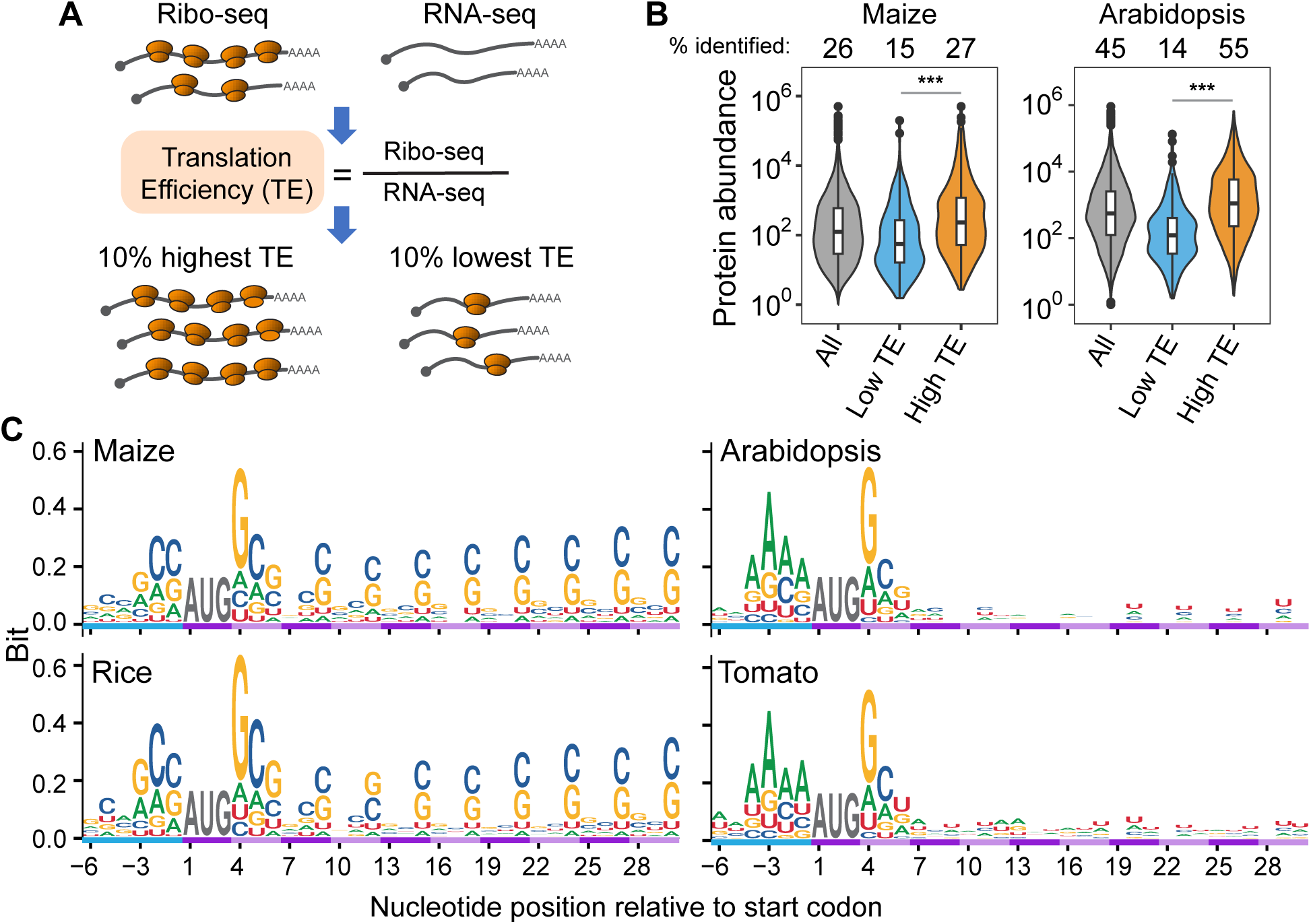
Comparison of high-TE and low-TE genes across four plant species. **A)** Workflow of identification of high-TE and low-TE genes. **B)** Violin plots showing high-TE genes are associated with higher protein abundance measured by quantitative proteomics. **C)** Sequence bias of high-TE genes in four model species. While all four species show strong Kozak sequence bias, enrichment of G and C in the 3^rd^ nucleotide of each codon (GC3) was only observed in monocots (maize and rice) but not in eudicots (Arabidopsis and tomato). Sequence logos displaying 6 nt upstream and 30 nt downstream from the CDS start codon are shown.

Given that 5′ UTRs can strongly influence translation initiation, we first performed a motif search using 5′ UTR plus 20 nt of coding sequences in high-TE genes, with sequences from low-TE genes serving as the control. While several motifs within the 5′ UTR were enriched in high-TE genes in various species, e.g., UA-rich and CA-rich regions in Arabidopsis, maize, and rice, as well as AGC motifs in maize and rice, we found the Kozak sequence was consistently identified as the most significant motif across all four species (**Supplemental Fig. 1**). Sequence logo comparisons confirmed that high-TE genes display a stronger bias in the Kozak sequence relative to low-TE genes (**Fig. 1C** and **Supplemental Fig. 2**), consistent with the established role of the Kozak sequence in regulating translation initiation ^45^. The monocots maize and rice prefer GCC**<u>AUG</u>**GCG, whereas the eudicots Arabidopsis and tomato prefer AAAA**<u>AUG</u>**GC, coinciding with the GC-rich genomes of maize and rice, in contrast to the AT-rich genomes of Arabidopsis and tomato.

Besides Kozak sequences, we observed that both maize and rice display sequence bias extending further into the coding sequence. Specifically, the 3^rd^ nucleotide of each codon shows a strong preference for G or C (GC3) in high-TE genes (**Fig. 1C**), suggesting that GC3 codons are associated with increased translation. In contrast, the two eudicots do not exhibit sequence bias beyond the Kozak sequence (**Fig. 1C**).

### GC3 codons are associated with high TE

To investigate the role of GC3 codons in TE regulation, we examined what codons are favored by high-TE genes across the four species compared to low-TE genes (**Fig. 2A**). In maize and rice, we observed strong codon preference for each amino acid in high-TE genes. Importantly, the codons favored by high-TE genes are all GC3 codons. The only GC3 codons that are not preferred by high-TE genes are found in six-fold degenerate amino acids (UUG for Leu, and AGG for Arg). In contrast, in Arabidopsis and tomato, the bias for GC3 codons is obscured. Some amino acids, like Ile, Lys, Phe, and Tyr, show a minor preference for the GC3 codons (**Fig. 2A**).

**Figure 2:**
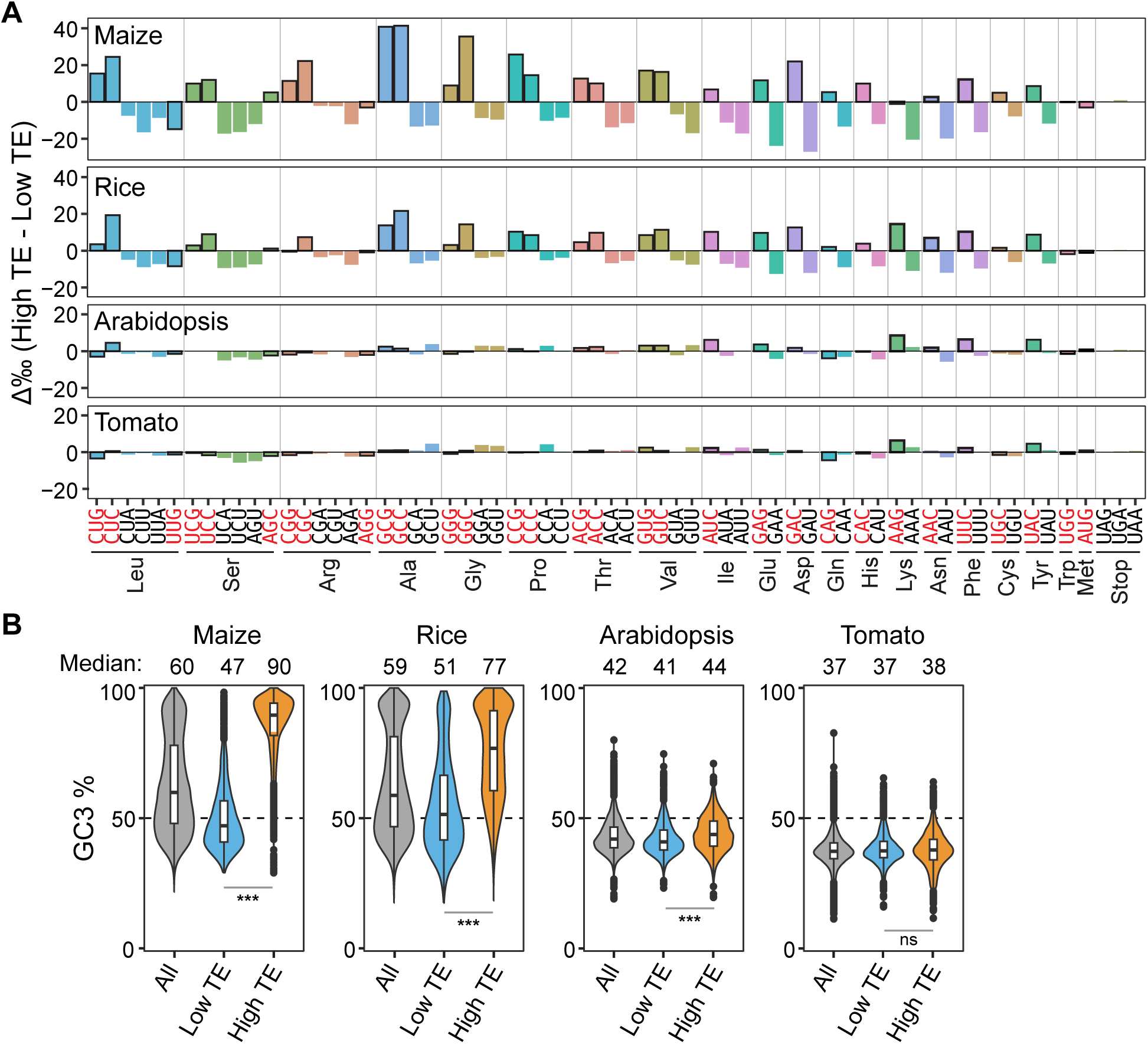
Higher TE is associated with increased use of GC3 codons. **A)** Differential codon usage of each amino acid between high- and low-TE genes. GC3 codons are highlighted in red font and outlined in black in the bar charts. **B)** Violin plots showing the distribution of GC3 content in all expressed genes, low-TE, and high-TE groups in the four species. Significance determined by Wilcoxon ranked sum test (ns: p > 0.05, *: p < 0.05, **: p < 0.01, ***: p < 0.001).

We next examined the overall CDS (coding sequence) GC3 content versus TE across species (**Fig. 2B**). In the GC-rich species maize and rice, we observed a striking difference in GC3 content between high-TE and low-TE genes, with high-TE genes having a median GC3 content of 90% and 77%, compared to 47% and 51% in low-TE genes, respectively (**Fig. 2B**). These results are consistent with previous rice polysome profiling data linking high GC3 with high TE, as well as with transgenic maize research showing that elevated GC3 increases target protein levels ^14,15^. In the AT-rich species Arabidopsis and tomato, AU3 codons are more common, so we expected that high-TE genes may have lower GC3. Surprisingly, high-TE genes in Arabidopsis still show a small but significant increase in GC3 compared to low-TE genes (median 44% vs. 41%) (**Fig. 2B**). In tomato, which has the lowest GC content of the four species, high-TE genes also exhibit slightly higher GC3, although the difference is not statistically significant (median 38% vs. 37%) (**Fig. 2B**).

To corroborate these findings, we analyzed an additional 22 Ribo-seq datasets, which produced consistent results across all four species from different growth conditions and tissue types (**Supplemental Table 1, Supplemental Fig. 3**). Together, these findings support that higher GC3 is correlated with higher TE, with much stronger differences observed in monocots compared to eudicots.

### Increasing GC3 improves protein production in both GC- and AT-rich species

To directly test whether high GC3 content can increase protein production in GC-rich and AT-rich species, we performed dual luciferase assays using luciferases with varying GC3 content but identical amino acid sequences. In the first set of plasmids, we designed Firefly luciferase (Fluc) with low (50%), mid (64%), and high (83%) GC3 content, and used Renilla luciferase (Rluc) as the internal control (**Fig. 3A**, **Supplemental File 1**). In maize, mid GC3 and high GC3 Fluc variants yielded 5.6-fold and 55-fold increase in relative luminescence (a proxy of Fluc protein levels) over the low GC3 (**Fig. 3C, left**). We observed a similar trend in Arabidopsis, with mid GC3 and high GC3 showing 2.2-fold and 4.1-fold higher luminescence than the low GC3 (**Fig. 3E, left**). In tobacco BY2 cells, which are in the same family as tomato, mid GC3 and high GC3 also showed 1.5-fold and 2.6-fold higher luminescence than low GC3 (**Fig. 3G, left**). These results demonstrate that high GC3 content can increase protein production, even in AT-rich species.

**Figure 3:**
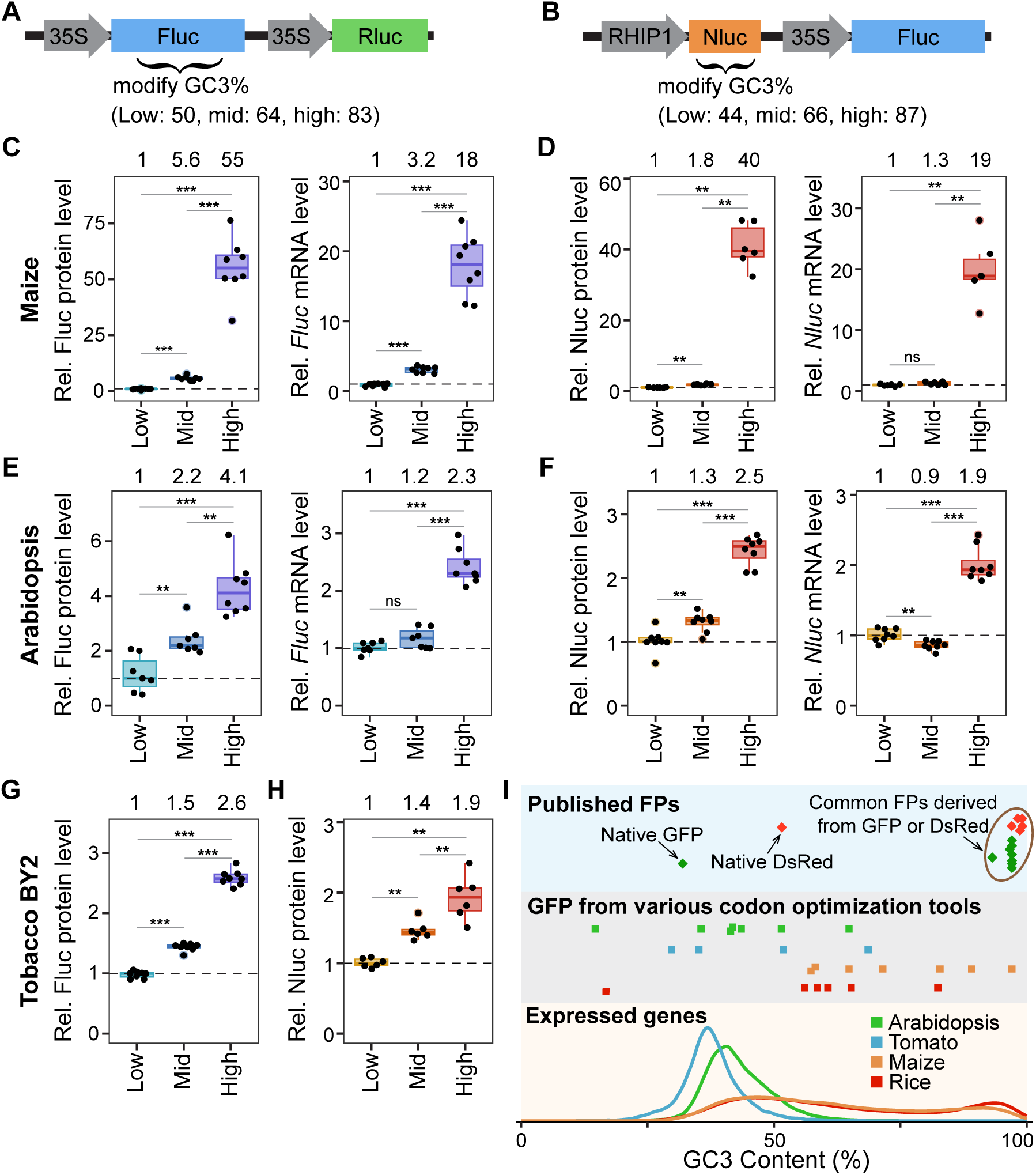
Higher GC3 codons increase protein production across plants, even for AT-rich species. **A-B)** Schematic of plasmids to test the effect of GC3 content on protein production using two different reporters. **C-H)** Relative protein levels from dual luciferase assay, relative mRNA levels from RT-pPCR quantified for Fluc (C, E, G) or Nluc (D, F, H) in maize (C, D), Arabidopsis (E, F), and tobacco BY2 protoplasts (G, H). The numbers above the graphs indicate the fold change of the median normalized to the low GC3 constructs. Significance determined by Wilcoxon ranked sum test (ns: p > 0.05, *: p < 0.05, **: p < 0.01, ***: p < 0.001). **I)** The commonly used fluorescent proteins (FPs) contain extremely high GC3. **Top:** Comparison of GC3 content of native GFP and DsRed versus widely used engineered versions. **Middle:** Suggested GC3 content for GFP using various codon optimization tools in Arabidopsis, tomato, maize, and rice. **Bottom:** GC3 content of expressed genes in the four model plants.

To validate this trend, we performed dual luciferase assays using a second set of plasmids, in which NanoLuc luciferase (Nluc) was modified to contain three levels of GC3 content (44%, 66%, and 87%), with Fluc serving as the internal control (**Fig. 3B**, **Supplemental File 1**). Consistently, the Nluc results demonstrated that higher GC3 content increased protein production in all three species (**Fig. 3D, F, left** and **3H**).

To determine whether the increase in Fluc and Nluc luminescence was driven by enhanced translation efficiency or greater mRNA abundance, we quantified the luciferase mRNA levels using RT-qPCR. The qPCR primers were designed within regions that are not codon-modified to ensure fair comparisons across different luciferase variants. Interestingly, in both maize and Arabidopsis, higher GC3 also increased *Fluc* and *Nluc* mRNA levels (**Fig. 3C-F, right**). However, the increase in mRNA levels alone cannot fully account for the observed rise in Fluc and Nluc protein levels, suggesting that enhanced translation efficiency by higher GC3 also contributes to increased protein production.

The increased Fluc and Nluc protein levels in Arabidopsis and tobacco contradict the conventional codon optimization guideline, which recommends matching the host genome’s codon frequency to boost protein production. The low GC3 Fluc and Nluc are the only two luciferases that fall within the normal range of GC3 content in Arabidopsis (**Supplemental Fig. 4**), yet these low GC3 variants consistently showed the lowest activity (**Fig. 3E, F, left** and **Fig. 3G, H**). Moreover, the low GC3 Nluc was specifically designed to match the codon usage of Arabidopsis high-TE genes, but it was still outperformed by both the original Nluc sequence (mid GC3) and the Nluc optimized to maize high-TE genes (high GC3) (**Fig. 3E, F**). Thus, increasing GC3 content represents a new promising strategy to enhance protein production in both AT- and GC-rich species.

To explore whether high GC3 content broadly enhances heterologous expression, we examined the GC3 composition of other commonly used reporter genes. Fluorescent proteins (FPs), such as GFP and DsRed, are widely used across many organisms and have been extensively engineered to improve expression and brightness ^46^. Compared to the native GFP and DsRed (32% and 52% GC3), the current commonly used variants show strikingly higher GC3 content (93–97% for GFP; 97–99% for DsRed) (**Fig. 3 I, top, Supplemental Table 2**). The extremely high GC3 content of these FPs well exceeds the GC3 levels recommended by various codon-optimization tools and most of the expressed genes in Arabidopsis, tomato, rice, and maize (**Fig. 3 I, middle** and **bottom**). The widespread adoption of high-GC3 FPs further supports the capacity of GC3 to enhance protein production across diverse species, regardless of their endogenous GC3 content.

### TE and GC3 are positively correlated across diverse lineages

We further investigated whether the preference for GC3 codons in high-TE genes is conserved across broader lineages. Because Ribo-seq data are only available for a limited number of species, we asked whether data from one species could be used to make predictions in another. To explore this, we first identified orthologs among the four model species using OrthoFinder ^47^, and then examined the correlation of TE between the orthologs in different species across different Ribo-seq datasets. We observed a consistent positive correlation in all species analyzed, with the strongest interspecies correlation between maize and rice (ρ = 0.52–0.58), and the weakest between rice and tomato (ρ = 0.27–0.50) (**Fig. 4A**). Orthologs of Arabidopsis low-TE and high-TE genes in tomato, maize, and rice showed a consistent and significant difference in TE, validating our approach for identifying low-TE and high-TE gene groups based on orthology (**Fig. 4B**). Likewise, using ortholog predictions reproduced similar findings that high-TE genes are associated with higher GC3 (**Fig. 4C**), supporting the applicability of this approach to species lacking Ribo-seq data.

**Figure 4:**
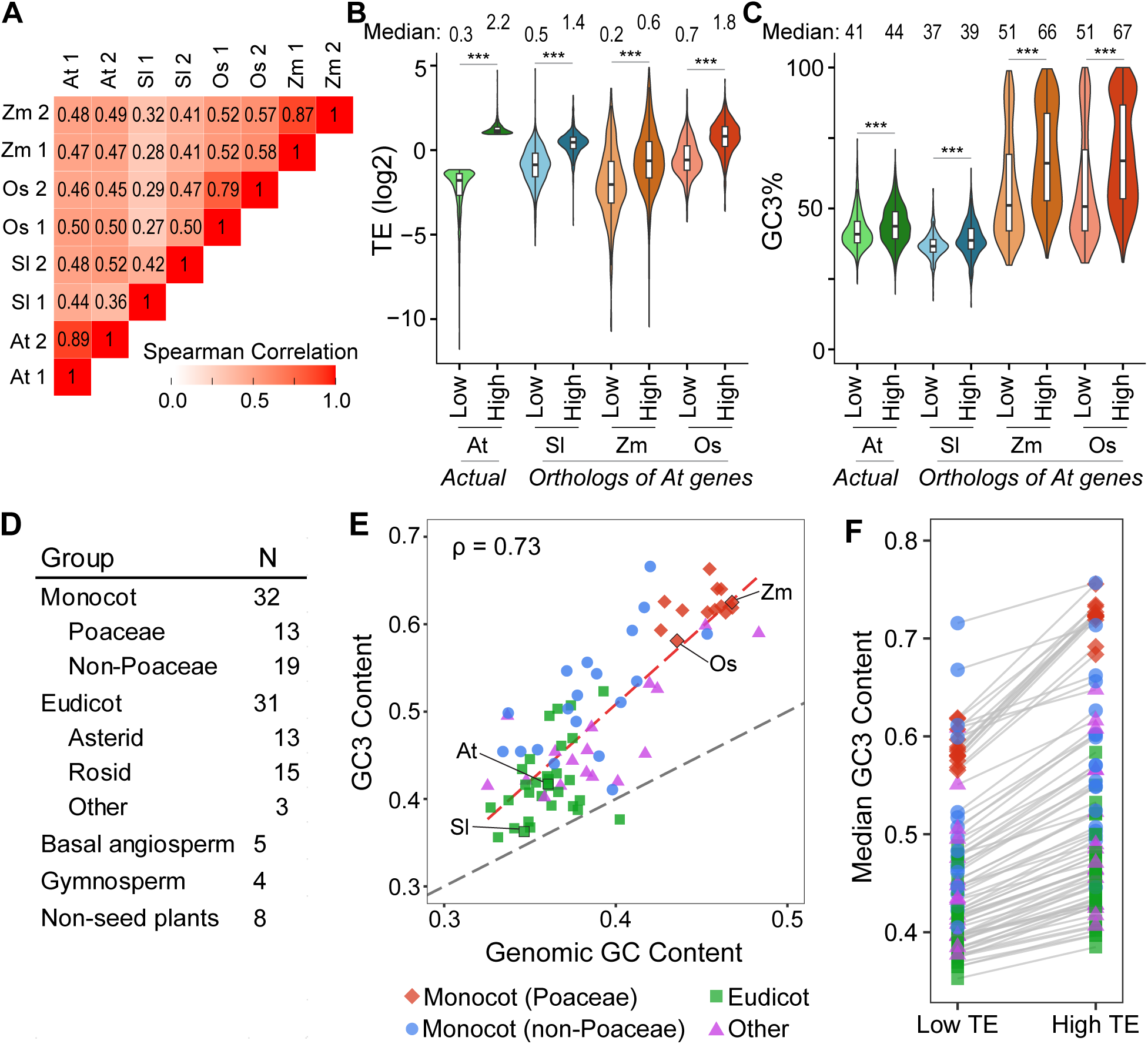
High TE is consistently associated with high GC3 across diverse species. **A)** Spearman correlation coefficients of TE between different Ribo-seq datasets from Arabidopsis (At), tomato (Sl), rice (Os), and maize (Zm), comparing the same genes if the data is from the same species, or orthologs if comparing two different species. **B-C)** Violin plots of TE (B) and GC3 (C) of Arabidopsis low-TE and high-TE genes, or of orthologs of Arabidopsis low-TE and high-TE genes in tomato, maize, and rice. Significance determined by Wilcoxon ranked sum test (ns: p > 0.05, *: p < 0.05, **: p < 0.01, ***: p < 0.001). **D)** Categorization of 80 species used in this analysis. **E)** Scatter plot showing that GC3 content is positively correlated with total genomic GC content in the 80 species. Red dashed line: the regression slope; grey dashed line: one-to-one relationship. **F)** Median GC3 content of low-TE and high-TE genes for each of the 80 species identified using orthologs of Arabidopsis low-TE and high-TE genes. Legend shared with (E). Grey lines connect medians for the same species.

We selected 80 diverse plant species, including a wide range of angiosperms, gymnosperms, and non-seed plants (**Fig. 4D, Supplemental Table 3**). Their GC3 content ranges from a minimum of 35.6% in the model legume (*Medicago truncatula*) to a maximum of 66.6% in the duckweed *Spirodela polyrhiza* (**Supplemental Table 3**). Most angiosperms outside of Poaceae, including early-diverged angiosperms, eudicots, and non-Poaceae monocots, have GC3 content well below that of the Poaceae cluster (**Fig. 4E**). Overall, GC3 content is strongly correlated with genome GC content across species (ρ = 0.73, **Fig. 4E**). Notably, the regression slope between genomic GC content and GC3 content (red dashed line) exceeds the one-to-one linear relationship (grey dashed line) (**Fig. 4E**), suggesting a selective bias favoring higher GC3 content.

We applied the ortholog-based TE prediction strategy to the 80 diverse species. In all species, the median GC3 content of the high-TE genes is higher than that of the low-TE genes, with statistically significant differences in all 80 species (**Fig. 4F**). Together, our ortholog-based TE predictions suggest that high TE is positively correlated with high GC3 across diverse species. This result is consistent with our observation that high GC3 increases heterologous protein production across both GC-rich and AT-rich species (**Fig. 3**).

### Endogenous high-abundance proteins exhibit increased GC3 across diverse lineages

We further investigated the relationship between GC3 and protein levels of endogenous genes. Mining Arabidopsis and maize proteomics data, we observed that high-protein genes have significantly higher GC3 content across species (**Fig. 5A, B**). This raised the possibility that high-protein genes may have evolved to adopt high GC3 content to enhance their expression throughout evolution. To address this, we identified homologs encoding three high-abundance protein groups across the 80 species: 1) Rubisco small subunit proteins (RBCS), 2) light-harvesting complex components (LHC), and 3) ribosomal proteins (RP) (**Supplemental Fig. 5** and **Supplemental Table 4**). We then compared their GC3 content relative to the species’ median GC3. Remarkably, all three protein groups displayed significantly higher GC3 content across a wide range of species (**Fig. 5C-F**). On average, the GC3 content of RBCS, LHC, and RP was 21%, 16%, and 10% above the species’ median GC3 across diverse lineages (**Fig. 5C-F**). The degree of increase in GC3 content also correlates with their protein abundance (**Supplemental Fig. 5**). These observations suggest that high GC3 is a conserved feature linked to the high protein abundance of these important genes across evolution.

**Figure 5:**
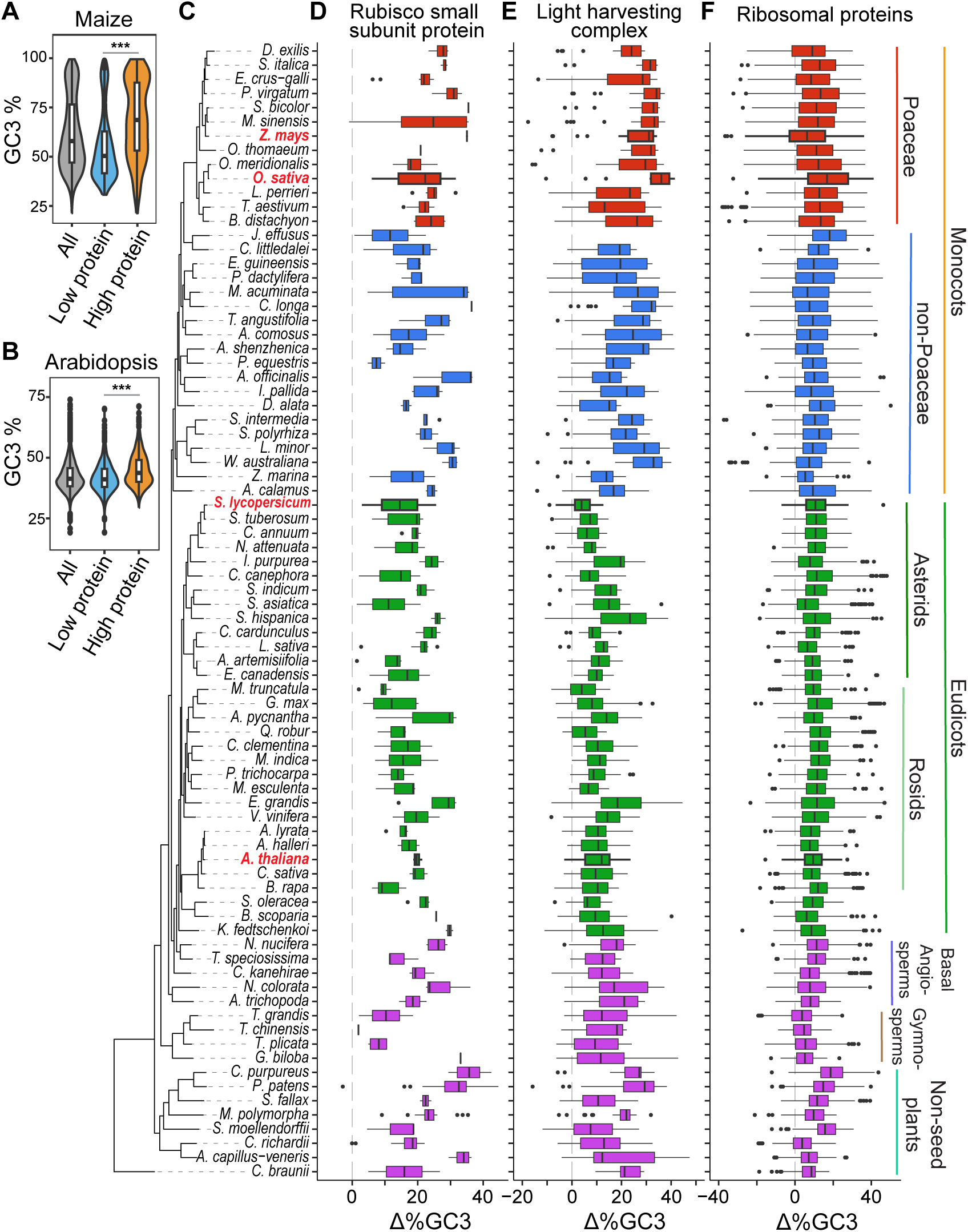
Important highly expressed proteins preferentially adopt high GC3. **A-B)** Violin plots showing that high-protein genes are associated with higher GC3 in maize and Arabidopsis (***: p < 0.001, Wilcoxon ranked sum test). **C)** Phylogenetic tree of 80 plant species from OrthoFinder. The four model species are highlighted in red. **D-F)** Boxplots illustrating differences in GC3 content between D) Rubisco small subunit (RBCS), E) light harvesting complex (LHC), and F) ribosomal protein (RP) genes, and the median GC3 content of all genes in each species. Homologs of Arabidopsis RBCS, LHC, and RP genes were used to identify these important, highly abundant protein genes. The grey vertical dashed line indicates 0 (no difference compared to the median of all genes).

### GC3 codons better match the tRNA pool across diverse species

To elucidate why higher GC3 broadly increases protein production, we tested the hypothesis that GC3 codons may better match the tRNA pool across diverse species. To evaluate this, we curated high-confidence tRNA genes in the 80 plant species with tRNAscan-SE ^48^ and quantified anticodon counts as a proxy for tRNA availability (**Supplemental Table 5**). The total number of tRNA genes identified ranged from 153 to 4,066, with a median of 577 (**Supplemental Fig. 6A**). The number of distinct anticodons identified in each species ranged from 45 to 53, with a median of 48 (**Supplemental Fig. 6B**). The anticodon counts across species consistently showed positive correlations, with more closely related species generally exhibiting stronger correlations (**Supplemental Fig. 6C**).

To connect tRNA anticodon availability to codon usage, we next calculated the codon weight (W_i_) for each codon, which quantifies codon optimality (ranging from 0 to 1, where 0 is nonoptimal and 1 is optimal) (**Supplemental Fig. 7, Supplemental Table 6**). W_i_ is determined by the corresponding tRNA abundance and anticodon-codon base pairing efficiency, with penalties applied for wobble base-pairing ^19^. We found that W_i_ values vary greatly across species, although some codons are consistently high, such as GAG for Glu (average 0.92), while others are consistently low, such as UUA for Leu (average 0.26) (**Supplemental Fig. 7**). Although the GC3 codons for each amino acid do not always have the highest W_i_ (**Supplemental Fig. 7**), on average, codons with a 3^rd^ position G or C have higher W_i_ than those ending with A or U (**Fig. 6A**).

**Figure 6:**
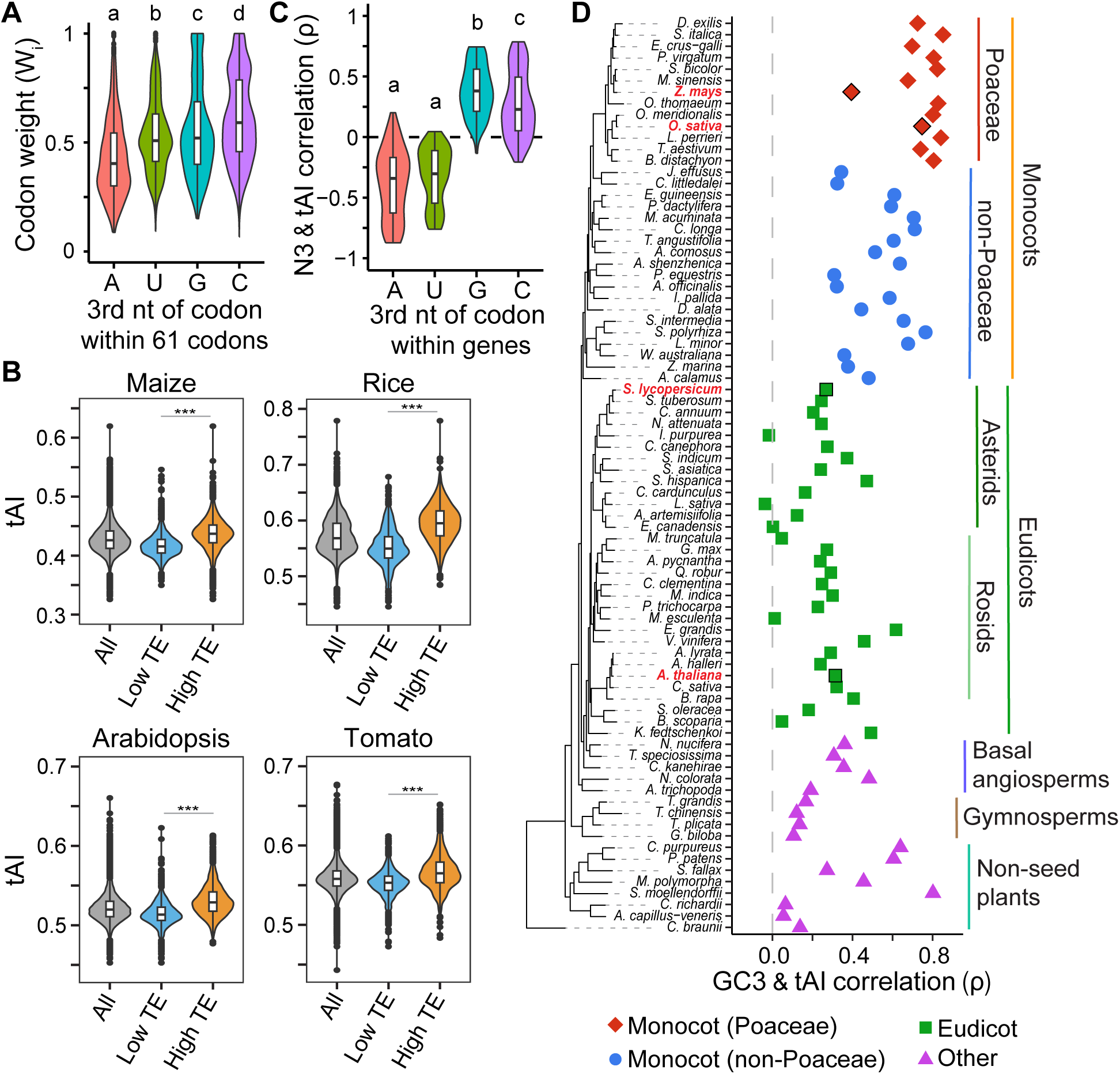
tRNA availability favors GC3 codons across different lineages. **A)** Codons with a third nucleotide of G or C have higher codon weight (Wi) on average. Violin plots show the distribution of Wi for all 61 sense codons across 80 plant species, grouped by the third nucleotide of each codon. Letters a–d represent significant differences as determined by ANOVA followed by Tukey’s HSD (p < 0.05). **B)** High tAI is associated with high TE across plants. Violin plots of tAI for all expressed genes, low-TE and high-TE genes for maize, rice, Arabidopsis, and tomato, are shown. Significance determined by Wilcoxon ranked sum test (ns: p > 0.05, *: p < 0.05, **: p < 0.01, ***: p < 0.001). **C)** Codons with a third nucleotide of G or C show stronger positive correlations with tAI compared to those ending with A or U. Violin plots display the distribution of correlations between tAI and the frequency of specific third-position nucleotides across all genes in the 80 species. **D)** GC3 content is positively correlated with tAI across diverse plant lineages. Spearman’s rank correlation (ρ) between GC3 and tAI is shown for the 80 species. The four model species are highlighted. The grey vertical dashed line indicates ρ =0.

To determine how all codons in a gene collectively match to the tRNA pool, we calculated the tRNA adaptation index (tAI), which incorporates W_i_ values for all codons in the CDS ^19^. In maize, rice, Arabidopsis, and tomato, tAI is positively associated with TE, as well as RNA-seq and Ribo-seq measurements (**Fig. 6B, Supplemental Fig. 8**). This suggests that genes better matching the tRNA pool also exhibit higher mRNA abundance and translation. Importantly, we found that tAI is almost always positively correlated with frequency of G3 and C3 codons within genes (**Fig. 6C**), as well as GC3 content overall (**Fig. 6D**), across the 80 plant species. Conversely, the frequencies of A3 and U3 codons are nearly always negatively correlated with tAI (**Fig. 6C**). This provides a mechanistic explanation for why higher GC3 can improve protein production across diverse plants, including many AT-rich species. Although eudicots generally have low GC3 content (**Fig. 4E**), their tAI values are still positively associated with GC3 (ρ = 0.24 on average, **Fig. 6D**). In contrast, lineages with higher GC3 content, such as Poaceae, show a much stronger correlation between GC3 and tAI (ρ = 0.75 on average, **Fig. 6D**). Together, the relationship between tAI and GC3 suggests that higher GC3 enhances protein production by better matching the tRNA pool, which also correlates with elevated mRNA abundance and translation efficiency.

### *GC3 Recoder*: a tool for customizing codon usage

To help researchers easily modify the GC3 content of a CDS, we developed an online app, *GC3 Recoder* (https://larrywu.shinyapps.io/GC3-recoder/) (**Supplemental Fig. 9**). Users can adjust GC3 content to a target percentage or match codon frequencies specified by a custom table, such as the maize high-TE gene codon frequency table used to generate the high-GC3 luciferases in this study, with the option to preserve specific restriction enzyme sites.

## DISCUSSION

Contrary to the conventional recommendations to match the codon frequency of the host genomes ^49^, our findings suggest a new codon optimization strategy: increasing GC3 content can enhance protein production across diverse plant species, including both GC- and AT-rich genomes. Because AT-rich species generally have a narrower GC content distribution, their genomes contain fewer high-GC3 genes; therefore, the effect of high GC3 is more difficult to detect. Yet important and highly abundant proteins, such as the Rubisco small subunit and ribosomal proteins, preferentially adopt high GC3 throughout evolution. In addition to our luciferase experiments, the widely used high-GC3 GFP and DsRed further highlight the applicability of GC3 for improving protein expression across diverse species.

High GC3 content is easy to achieve, as every amino acid has at least one GC3 codon, making it theoretically possible to encode any protein with 100% GC3 codons. In our case, we simply matched the codon usage of maize high-TE genes. We also developed an online tool, *GC3 Recoder*, to help researchers easily modify the GC3 content for gene engineering.

Our results, together with previous findings, strongly suggest that high GC3 content synergistically enhances protein production through multi-level regulatory mechanisms. In addition to boosting translation efficiency and mRNA abundance, high GC3 may also reduce DNA methylation and small RNA-mediated gene silencing, as well as disrupt AU-rich motifs that destabilize mRNAs ^15,50,51^. Our results further suggest that GC3 codons better match tRNA availability. One possible reason for the universally GC3-favoring tRNA pools in diverse species is that GC base pairing enables stronger codon–anticodon hydrogen bonding, which could promote faster and more efficient translation ^17,52^.

Beyond plants, high GC3 in humans also increases mRNA level and protein production, both in reporter assays and in endogenous genes ^53,54^. Numerous species, including many animals and fungi, have a large tRNA repertoire, where matching tRNAs are available for GC3 codons. Thus, provided the corresponding anticodons are present, GC3 preference may be generalized to other eukaryotes.

## MATERIALS AND METHODS

### Ribo-seq and RNA-seq processing

The code and parameters used are provided in the accompanying GitHub (https://github.com/kaufm202/GC3_codon_use). Briefly, raw Ribo-seq and RNA-seq data were downloaded from NCBI Short Read Archive (SRA) or Amazon Web Services (AWS) using the accession IDs in **Supplemental Table 1**. Adapter and quality trimming were performed using cutadapt ^55^. rRNA and tRNA sequences were acquired from the NCBI Nucleotide database and removed using Bowtie2 ^56^. The filtered reads were then aligned to CDS from the Araport 11 annotation for Arabidopsis, the MSU release 7 annotation for rice, the iTAG v5.0 annotation for tomato, and the NAM 5.0 annotation for maize using STAR ^57^. RSEM was then used to estimate transcripts per million (TPM) from the STAR alignments ^58^. Translation efficiency (TE) was calculated as the mean Ribo-seq TPM divided by the mean RNA-seq TPM within CDS for each transcript. Only the most abundant transcript isoform from each gene and transcripts with RNA-seq TPM > 1 and TE < 30 were used for downstream analysis. The genes with the 10% lowest and highest TE were defined as the low-TE and high-TE genes in each species.

### Analysis of quantitative proteomics data

Quantitative proteomics data from label-free mass spectrometry was downloaded from published datasets for Arabidopsis ^59^ and maize ^60^, using label-free quantification (LFQ) values in Arabidopsis, and distributed normalized spectral abundance factor (dNSAF) in maize. To match the published dataset, the maize Ribo-seq and RNA-seq were realigned to the B73 RefGen_v2 5a genome annotation. Abundance values within each species were normalized to the minimum value.

### Motif analysis

The 5′ UTR plus 20 nt of the CDS of each transcript was extracted using the GenomicFeatures package ^61^ in R ^62^. For transcripts without annotated 5′ UTRs, the 100 nt upstream of the CDS start codon were assumed as the 5’UTR. Motif discovery in high-TE genes was performed using STREME from the MEME suite ^63,64^ using the low-TE genes as the control with a p-value cutoff of 0.05.

### Modification of GC3 content of Fluc and Nluc and construction of dual luciferase assay (DLA) plasmids

The sequences of different Fluc and Nluc luciferase variants with different GC3 contents are provided in **Supplemental File 1**. Fluc and Nluc were codon-modified to modulate GC3 content without changing the peptide sequence using the ExpOptimizer codon optimization tool available online through NovoPro (www.novoprolabs.com/tools/codon-optimization). For the Fluc sequences, the first 20 codons were unmodified to avoid potential effects from the Kozak sequence, then the following 372 codons were modified, with PpuMI, KasI, and BamHI RE sites excluded. The Low GC Fluc contains the unmodified Fluc sequence of the pHsu133 plasmid ^9^ (see below). The Mid GC Fluc was codon-modified with the default codon usage table provided by NovoPro for *Zea mays* as the expression host. The High GC Fluc was codon optimized with a user-defined codon usage table derived from our maize high-TE genes (**Supplemental File 2**). For Nluc, the first 10 codons, qPCR primer annealing sites, and HA tag sequence were unmodified, and the remaining 153 codons were codon-modified, with XbaI, KpnI, and EcoRI RE sites excluded. The Mid GC Nluc contains the original Nluc sequence of the pHsu510 plasmid (see below). The Low GC Nluc was codon-modified with a user-defined codon usage table derived from our Arabidopsis high-TE genes (**Supplemental File 2**). The High GC Nluc was codon-modified with a user-defined codon usage table derived from our maize high-TE genes (**Supplemental File 2**).

For the Fluc/Rluc DLA constructs, the pHsu133 plasmid ^9^ containing 35S-Fluc and 35S-Rluc was used as the Low GC reporter. The codon-modified Fluc variants were synthesized by BioBasic and cloned into pHsu133 via Kas I and PpuM I sites, generating the Mid GC and High GC Fluc plasmids.

For the Nluc/Fluc constructs, plasmid 3698 (a gift from Yves Poirier’s Lab) was used to generate the pHsu510 plasmid containing RHIP1-Nluc and 35S-Fluc, and used as the Mid GC Nluc plasmid. The codon-modified Nluc sequences were subcloned into pHsu510 via XbaI and KpnI sites, generating the Low GC and High GC Nluc plasmids.

### Protoplast transformation and dual luciferase assays

Protoplast isolation, transformation, and dual luciferase assays were performed as previously described ^9,65^. For Arabidopsis protoplasts, surface-sterilized Arabidopsis Col-0 seeds were cold stratified in water for 2 days at 4°C, then planted on soil and grown for 19-21 days with 16-hour light (∼100 μmol m^−2^ s^−1^, cool white, fluorescent bulbs) and 8-hour dark cycle at 22°C with intermittent watering. Fully expanded rosette leaves were excised and thinly sliced for protoplast isolation. Leaf slices were placed in enzyme solution (1% [w/v] cellulase, 0.25% [w/v] macerozyme, 0.4 M mannitol, 20 mM KCl, 20 mM MES, and 10 mM CaCl_2_), then vacuum infiltrated for 30 minutes, followed by a 2-hour incubation with gentle shaking at 40 rpm, and then 5 minutes at 80 rpm to release protoplasts. The protoplasts were filtered through a 70 μm cell strainer, then centrifuged at 100 x g at 4°C, and washed twice with cold W5 (154 mM NaCl, 125 mM CaCl_2_, 5 mM KCl, and 2 mM MES). The protoplasts were counted using a hemocytometer, then resuspended in cold MMG (4.5 mM MES [pH 5.7], 0.4 M mannitol, and 15 mM MgCl_2_) to a concentration of 1×10^5^ protoplasts per 150 μL.

For maize protoplasts, surface-sterilized maize B73 seeds were imbibed in water overnight at room temperature, then transferred to a petri dish with filter paper moistened with water for 3-4 days until germinated, then transferred to soil and grown at 22°C in the dark with intermittent watering. Leaves from 14-day etiolated maize seedlings were excised and thinly sliced ^66^. Protoplast isolation was then carried out with the same method as Arabidopsis, except with a 4-hour incubation in the enzyme solution.

For tobacco BY2 cell protoplasts, the cells were spread on petri dishes containing solid BY2 growth media (LS media plus 3% [w/v] sucrose, 0.56 mM myo-inositol, 3.8 µM thiamine, 4.4 mM K_2_HPO_4_, and 0.8% [w/v] Phytoblend agar) and grown for 10 days at room temperature in the dark. BY2 cells were then scraped directly from the plate for protoplast isolation. Protoplast isolation was carried out with the same method as for Arabidopsis, except with no vacuum infiltration, with a 3-hour incubation in enzyme solution, and using a 100 μm cell strainer at the filtering step.

6 to 8 replicates of 1×10^5^ protoplasts were mixed with 5 μg of plasmid DNA and 170 μL PEG solution (40% [w/v] PEG4000, 0.2 M mannitol, 100 mM CaCl_2_) and incubated for 5 minutes. Protoplasts were washed four times with cold W5, then transferred to cell culture plates and incubated in the dark for 16 to 18 hours. Protoplasts were then centrifuged at 2,250 x g for 3 min at 4 °C, the supernatant was removed, and then protoplasts were lysed by resuspending in 100 μL 1x Passive Lysis Buffer (Promega E1910). After shaking at room temperature for 15 minutes, protoplasts were centrifuged at 2,250 x g for 3 min at 4 °C, and 20 μL of cleared lysate was used for the luciferase assays. Fluc/Rluc and Nluc/Fluc DLAs were performed using the Dual-Luciferase Reporter Assay System (Promega E1960) and Nano-Glo Dual-Luciferase Reporter Assay System (Promega N1630), respectively, in accordance with the manufacturer’s protocols using the GloMax Navigator Plate Reader (Promega, GM2010). For Fluc/Rluc DLA, Fluc relative luminescence is reported as the Fluc luminescence normalized to Rluc luminescence for each sample, and then normalized to the median of the low GC samples. For Nluc/Fluc DLA, Nluc relative luminescence is reported as Nluc luminescence normalized to Fluc luminescence for each sample, and then normalized to the median of the low GC samples.

### RT-qPCR of DLA lysate

RNA was extracted from 60 µL DLA lysate using RNA Clean and Concentrator-5 kit (Zymo Research, R1016). To remove remaining DNA, purified RNA was then treated with 5 U of DNase I (Zymo Research, E1012) for 30 min, then repurified using RNA Clean and Concentrator-5 kit. cDNA synthesis was performed from 4 µL of eluted RNA using LunaScript RT SuperMix Kit (NEB, E3010) with a total reaction volume of 8 µL. qPCR was then performed using Luna Universal qPCR Master Mix (NEB, M3003) using primers provided in **Supplemental File 1**. Relative *Fluc* expression was calculated using the ΔΔCt method ^67^ using *Rluc* as the reference gene for Fluc/Rluc DLA, and relative *Nluc* expression was calculated using *Fluc* as the reference gene for Nluc/Fluc DLA, and normalized to the median of the low GC samples.

### GC3 Recoder online app

*GC3 Recoder* (https://larrywu.shinyapps.io/GC3-recoder/), a Shiny app for codon optimization, was developed to modify the input CDS codons using synonymous codons without altering the protein sequence. After checking the input sequence for errors and locking specific restriction sites, the app changes codons using one of two distinct methods specified by users. With the ‘Target GC3’ method, the app randomly selects synonymous codons to achieve the target GC3 content. With the ‘Frequency’ method, it selects synonymous codons based on a user-provided frequency table (consisting of 3 columns: codon, corresponding amino acid, and frequency per thousand).

### Phylogenetic analysis

Genome sequences and annotations for 80 species were downloaded from the sources described in **Supplemental Table 3**. CDSs were extracted and then translated to peptide sequences using GenomicFeatures ^61^ and BioStrings ^68^ in R ^62^. Only one CDS per gene was used, either the longest CDS when using all genes, or the CDS with the highest homology when using orthologs and homologs. Orthologs were identified using OrthoFinder ^47^. Arabidopsis Rubisco small subunit (RBCS), light-harvesting complex (LHC) and ribosomal protein (RP) genes were obtained from TAIR ^69,70^ and then removed genes with Ribo-seq TPM > 500. For homolog identification of the Arabidopsis RBSC, LHC, and RP genes, protein sequence were aligned using BLASTp with Diamond ^71^. We defined homologs as any protein with greater than 40% homology with at least one of the Arabidopsis proteins. The transcript IDs for the orthologs of Arabidopsis low-TE and high-TE genes and the homologs of the Arabidopsis RBCS, LHC, and RP genes in all 80 species are provided in **Supplemental Table 4**.

### Calculation of tAI

tRNA gene copies were identified using tRNAscan-SE ^48^. tRNA gene copies were filtered to a high confidence set by removing low-quality hits, pseudo-tRNAs, tRNAs with ambiguous anticodon, mitochondrial and plastidial tRNAs, and tRNAs with a high number of copies within a small genomic region. The total tRNA gene copies for each anticodon were then summed.

tRNA adaptation index (tAI) was calculated as previously described, with a minor modification to prevent outlier anticodon counts from skewing the final tAI values ^19^. Outliers were identified if the anticodon counts were greater than two standard deviations from the mean when the maximum anticodon count is excluded. Outlier anticodon counts were reduced to be equal to the next highest anticodon count within two standard deviations of the mean. Then, for each codon, the weight was calculated from the sum of anticodon counts for anticodons that pair with a given codon, with penalties for wobble base-pairing. The final codon weights (W_i_) represent the codon weight normalized to the codon with the maximum weight. tAI was calculated as the geometric mean of W_i_ of all codons in a given CDS, excluding the start and stop codons.

### Data visualization

Sequence logos were visualized using the ggseqlogo package ^72^ in R. Phylogenetic trees were visualized using ggtree ^73^. All other plots were generated using ggplot2 ^74^ in R.

### Code availability

The Bash and R scripts used to process the data as described above are available in the GitHub repository: https://github.com/kaufm202/GC3_codon_use.

## ACKNOWLEDGEMENTS

We thank Ning Jiang for the helpful discussion on the research. This work used computational resources and services provided by the Institute for Cyber-Enabled Research at Michigan State University. This work was supported by a predoctoral training award under Grant Number T32-GM110523 from the National Institute of General Medical Sciences of the National Institutes of Health to IDK, and research grants from the National Science Foundation under Award Numbers 2425390 and 2051885, and the National Institute of General Medical Sciences of the National Institutes of Health under Award Number R35GM155375 to PYH. The content is solely the responsibility of the authors and does not necessarily represent the official views of the NSF or NIH.

## AUTHOR CONTRIBUTIONS

IDK, HLW, and PYH designed the research and interpreted the data. IDK performed the analysis with input from HLW. IDK performed all experiments. HLW developed the *GC3 Recoder* Shiny app. IDK and PYH wrote the manuscript with input from HLW.

## SUPPLEMENTARY FIGURES

**Supplementary Figure 1:**
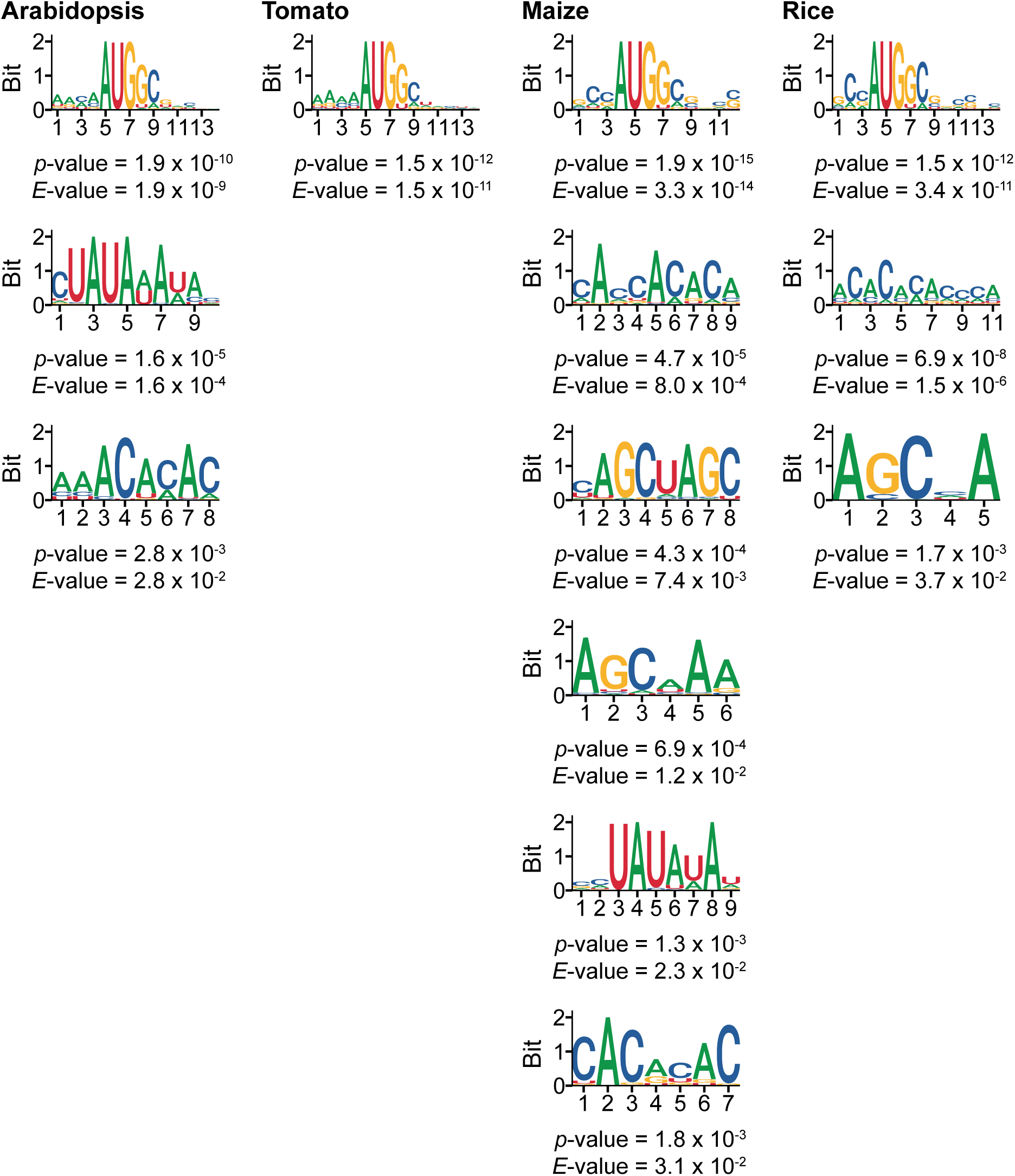
Motifs enriched in high-TE genes within the 5′ UTR plus 20 nt of the CDS region in Arabidopsis, tomato, maize, and rice. Motifs with *E*-value <0.05 are shown.

**Supplementary Figure 2:**
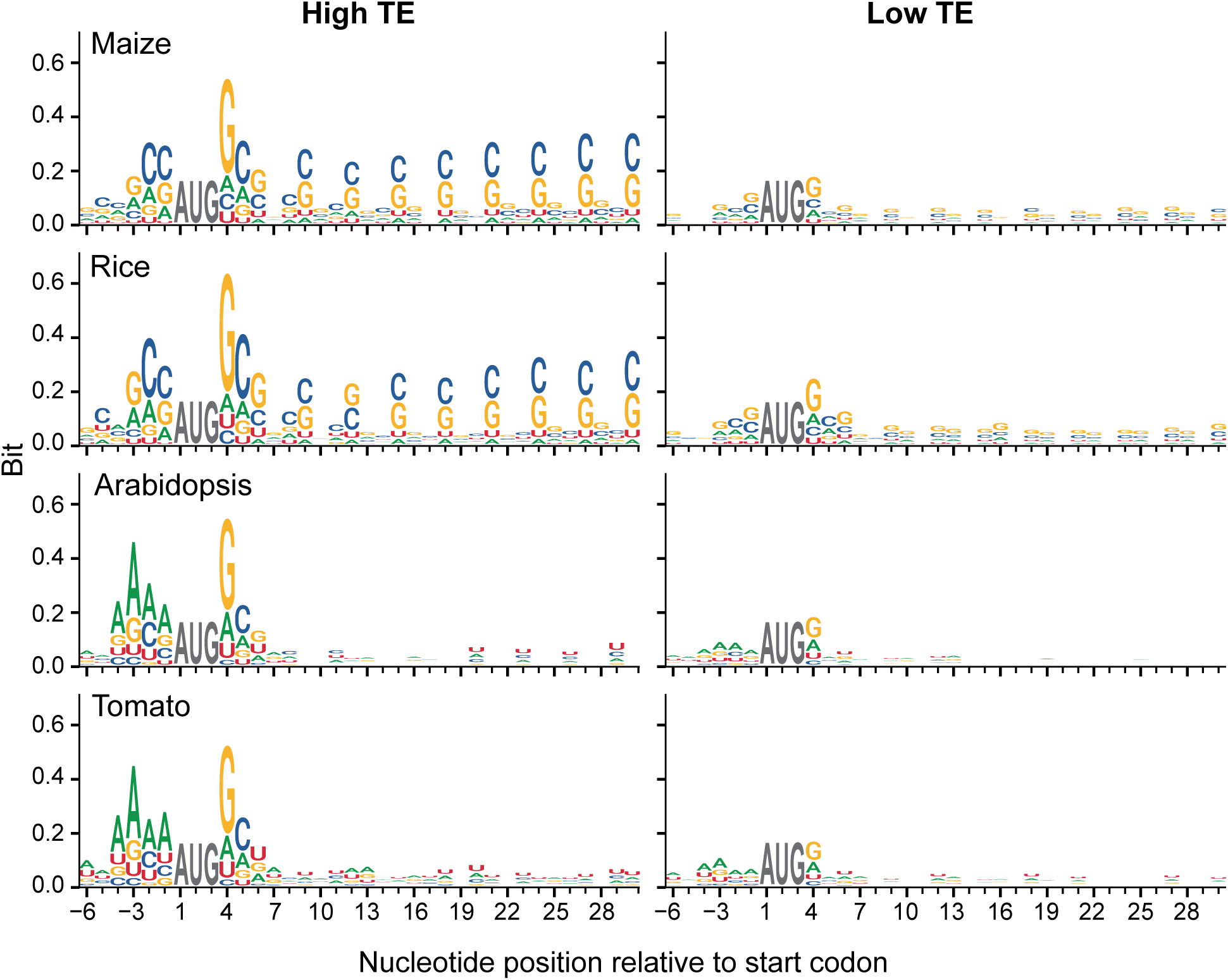
Sequence bias of high-TE genes (left) compared to low-TE genes (right). In contrast to strong Kozak in all four species, enrichment of G or C in the 3^rd^ nucleotide of each codon (GC3) was only observed in monocots (maize and rice) but not in eudicots (Arabidopsis and tomato). Sequence logos displaying 6 nt upstream and 30 nt downstream from the CDS start codon are shown.

**Supplementary Figure 3:**
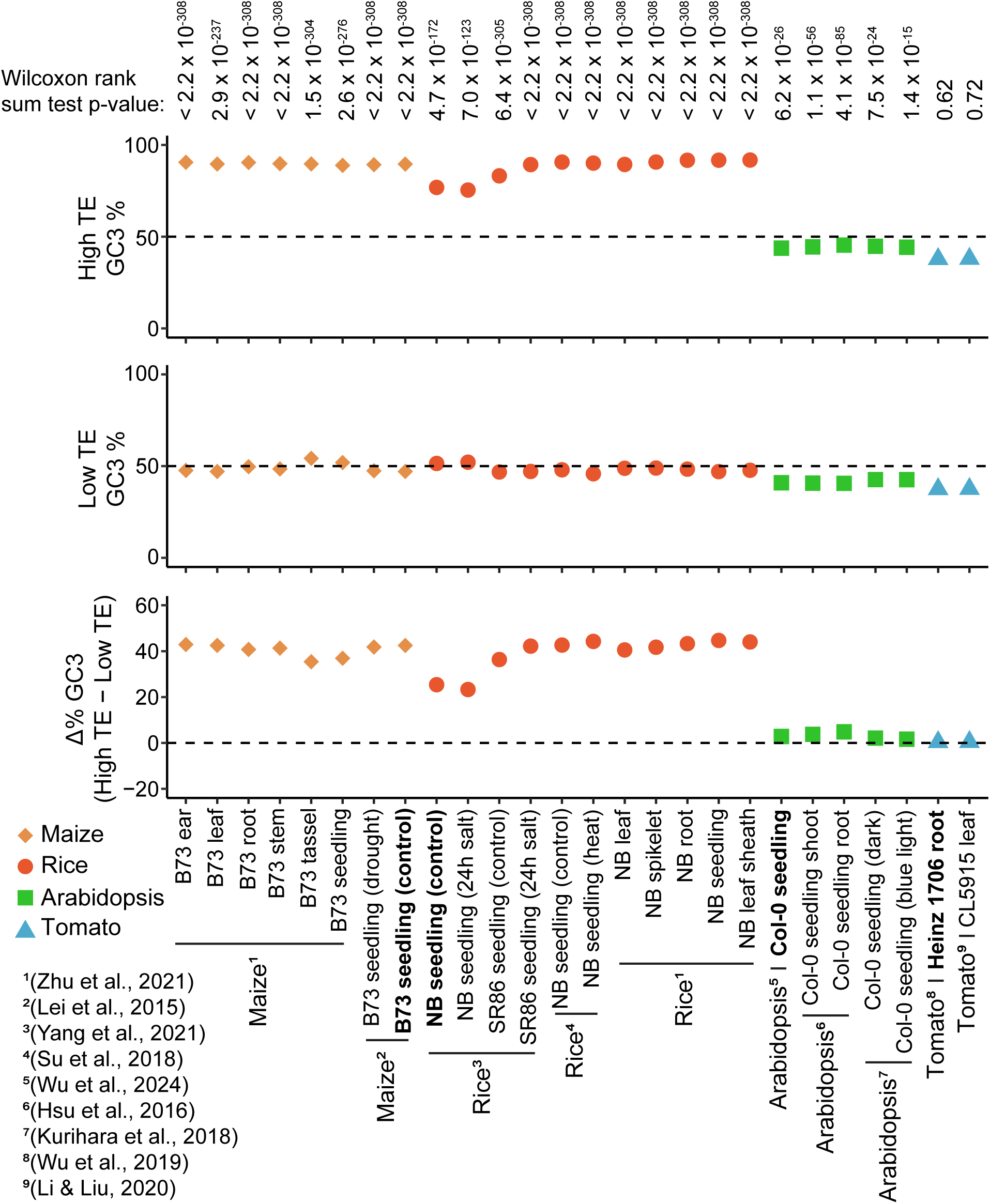
Median GC3 content of high-TE and low-TE genes, and the difference in GC3 content between high-TE and low-TE genes, across various tissues and conditions, are shown. Samples in bold are the samples presented in Fig. 2B. Ribo-seq data were previously reported (**Supplemental Table 1**).

**Supplementary Figure 4:**
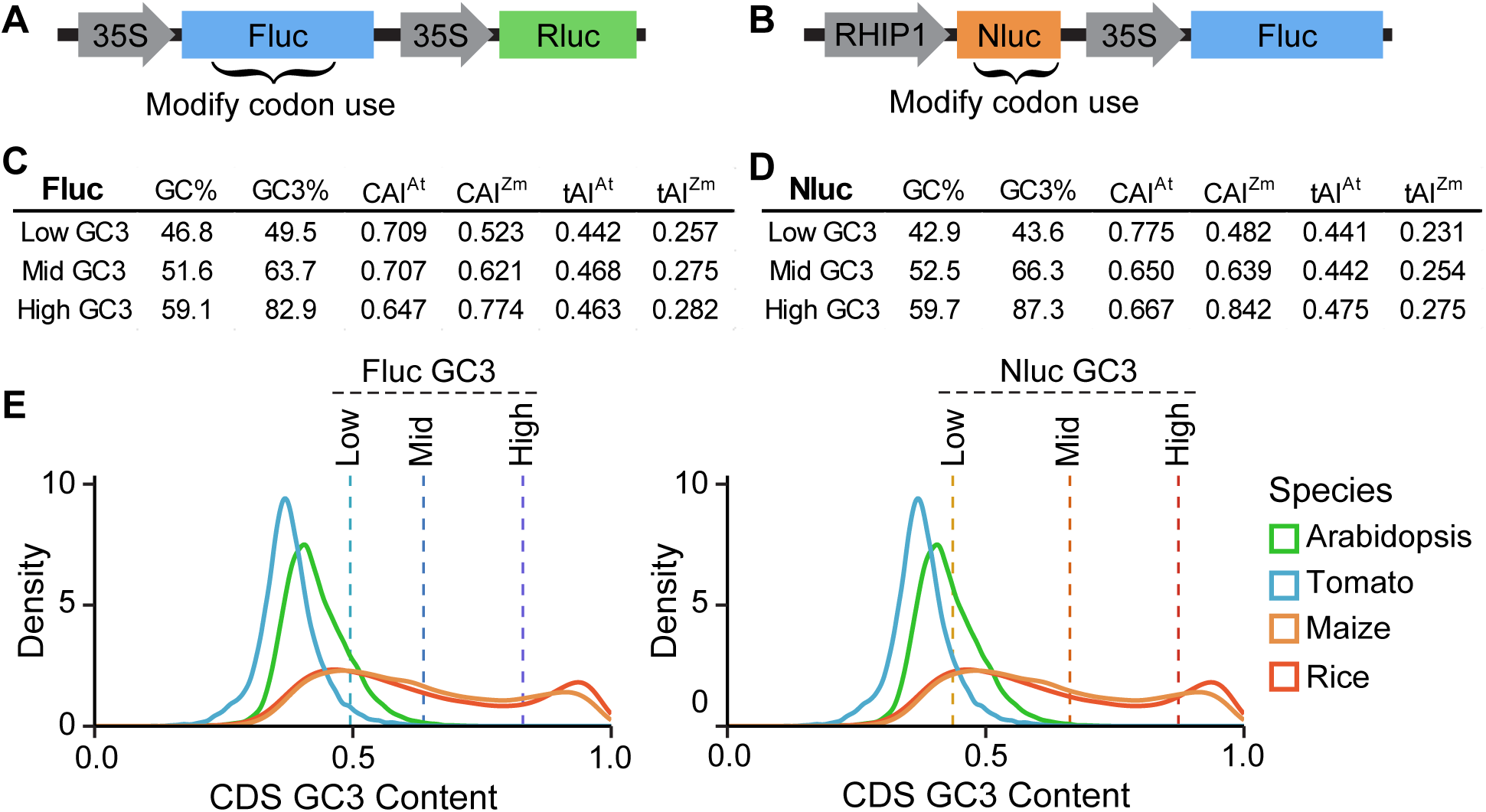
Additional information for Fluc and Nluc variants tested. **A-B)** Schematic of the **A)** Fluc/Rluc DLA and **B)** Nluc/Fluc DLA plasmids. **C-D)** Sequence content information for **C)** Fluc and **D)** Nluc variants. **E)** GC3 content distributions of expressed genes in Arabidopsis, tomato, maize, and rice, compared to the GC3 content of Fluc and Nluc variants.

**Supplementary Figure 5:**
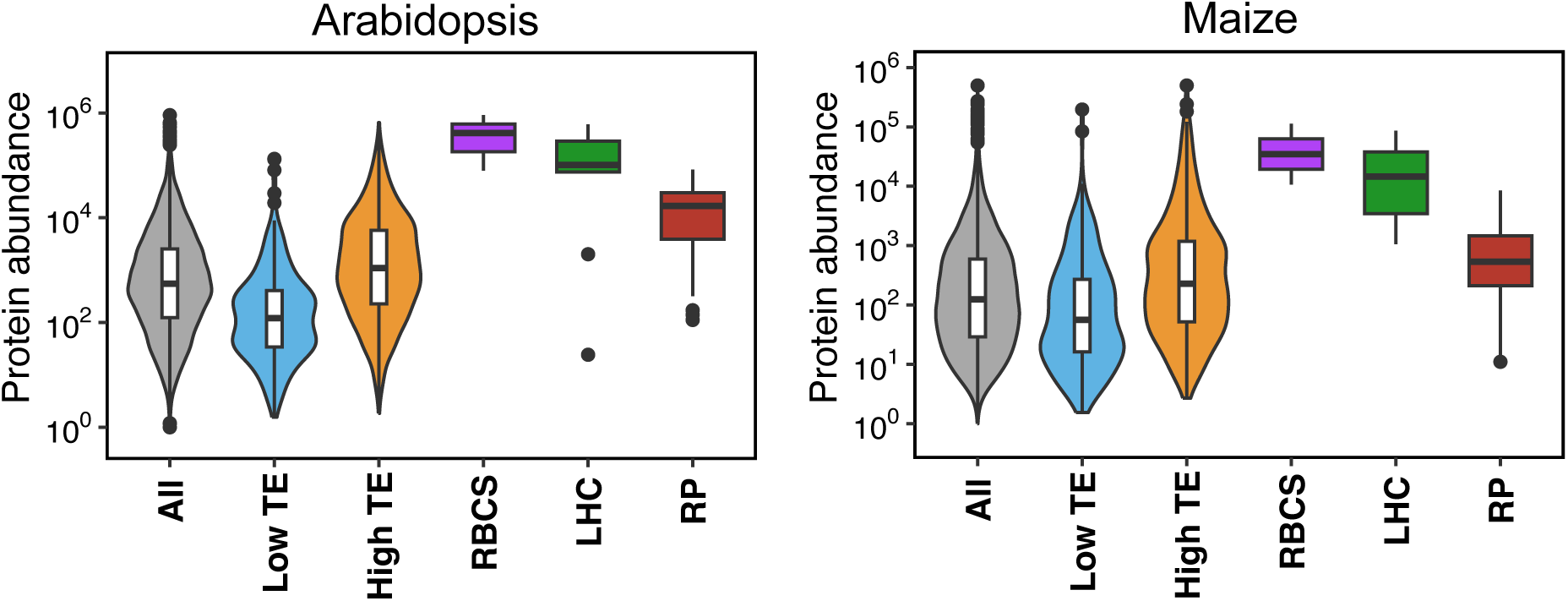
Protein abundance, as determined by quantitative proteomics, of Rubisco small subunit protein (RBCS), light-harvesting complex (LHC), and ribosomal proteins (RP) compared to other genes in Arabidopsis and maize.

**Supplementary Figure 6:**
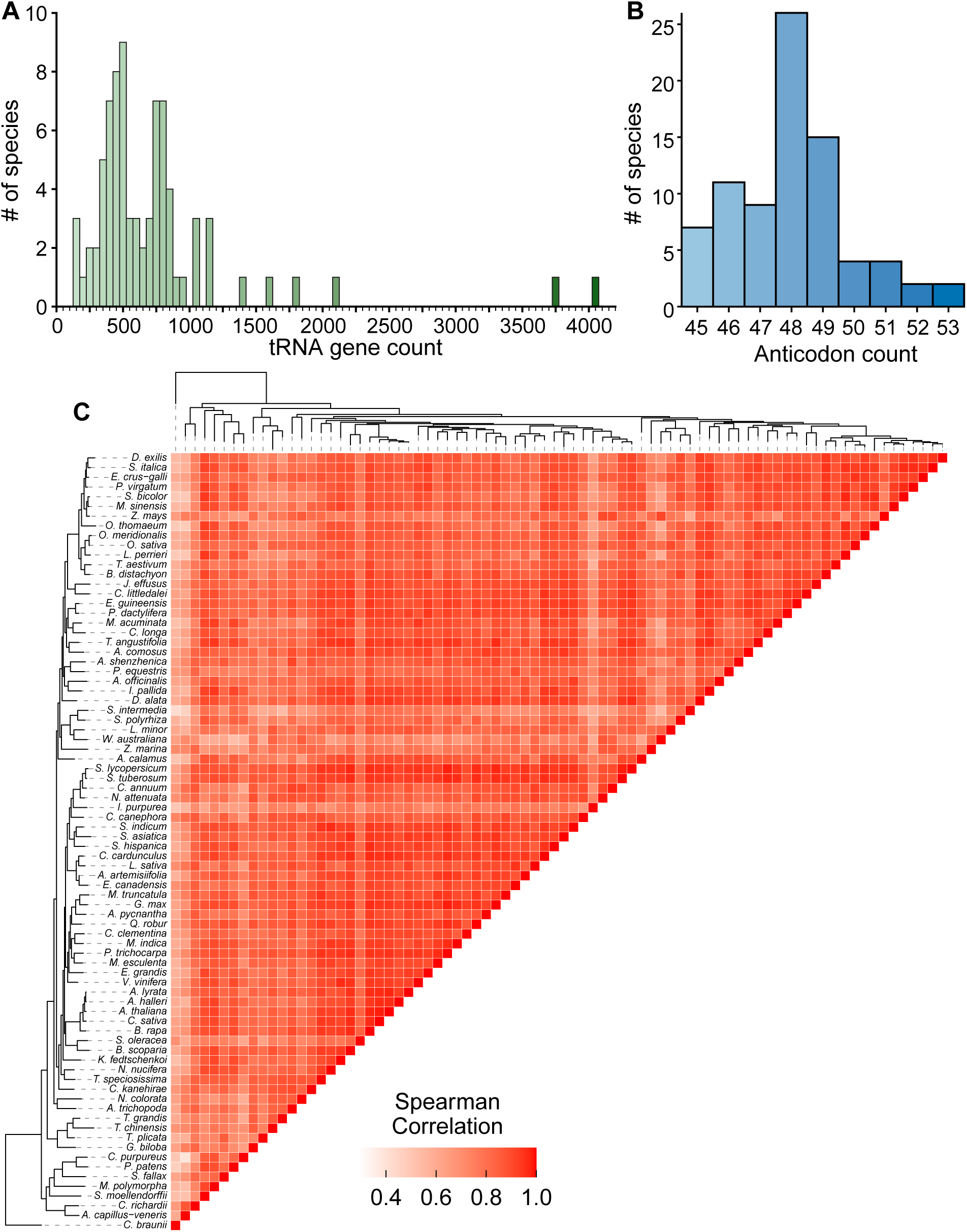
**A)** Histogram showing the distribution of tRNA gene count across the 80 plant species (bin = 50). **B)** Histogram showing the number of anticodons identified across the species. **C)** Correlation matrix of tRNA anticodon counts in the 80 plant species.

**Supplementary Figure 7:**
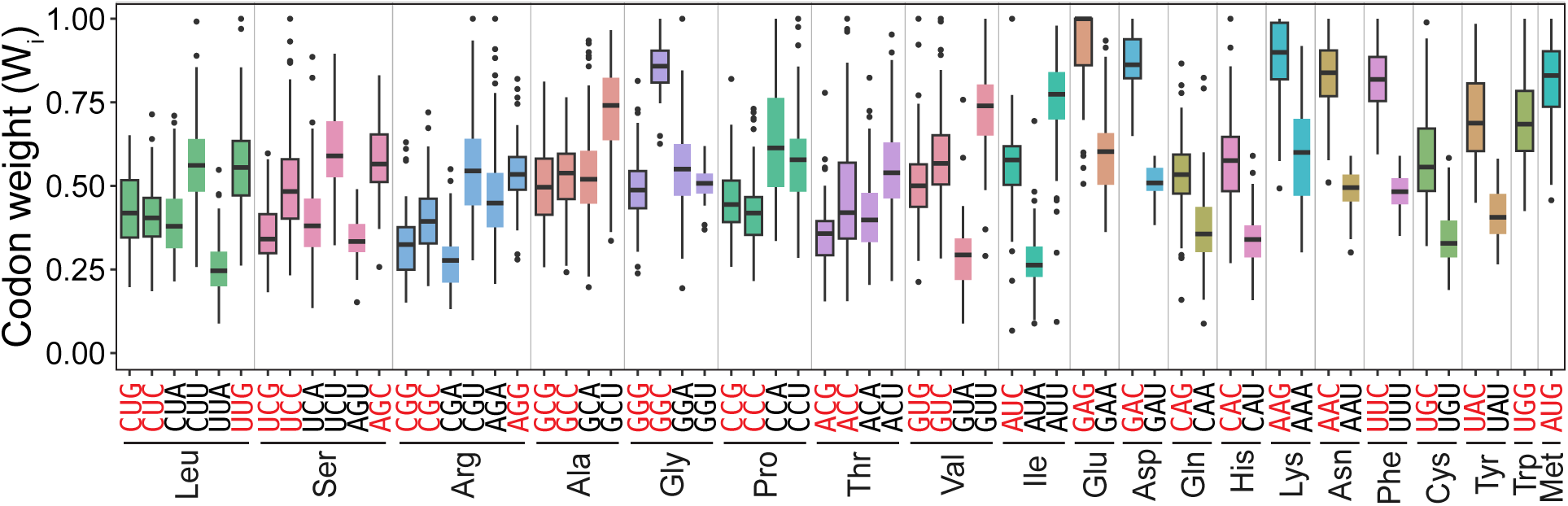
Box plots of codon weights (W_i_) for all 61 sense codons from 80 species. The GC3 codons for each amino acid are highlighted in red font and outlined in black in the box plots.

**Supplementary Figure 8:**
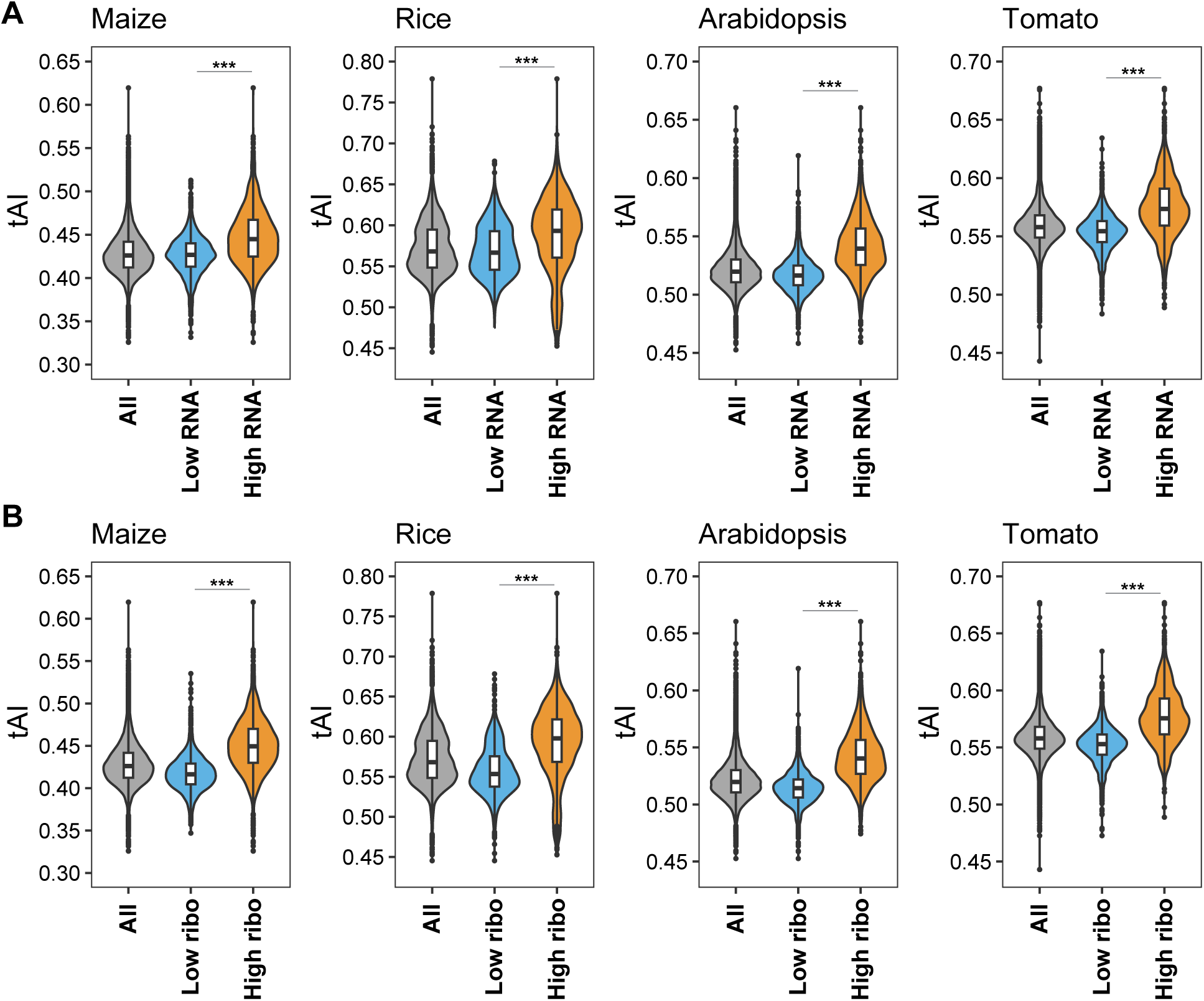
Relationship between tAI and RNA or translation levels. **A)** tAI values for all, low- and high-abundance RNAs determined by RNA-seq, and **B)** tAI values for all, lowly and highly translated RNAs determined by Ribo-seq in maize, rice, Arabidopsis, and tomato. Significance determined by Wilcoxon ranked sum test (ns: p > 0.05, *: p < 0.05, **: p < 0.01, ***: p < 0.001).

**Supplementary Figure 9:**
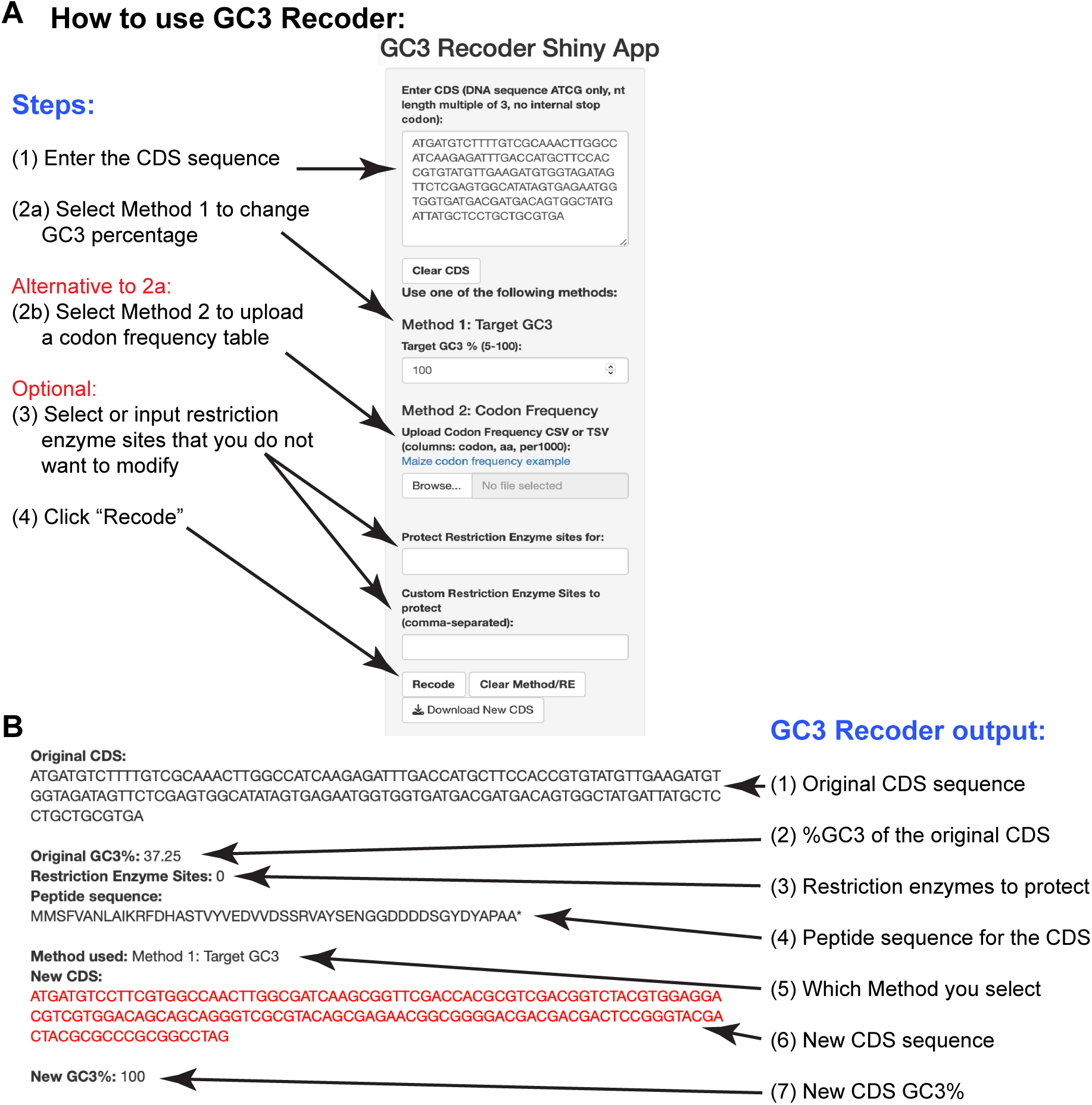
The *GC3 Recoder* Shiny App helps researchers modify codon usage with easy steps and flexible options. **A)** Steps for using *GC3 Recoder* to change synonymous codons based on target GC3 % or a user-defined codon frequency table. **B)** Output of *GC3 Recoder*.

